# multiGSEA: A GSEA-based pathway enrichment analysis for multi-omics data

**DOI:** 10.1101/2020.07.17.208215

**Authors:** Sebastian Canzler, Jörg Hackermüller

## Abstract

Gaining biological insights into molecular responses to treatments or diseases from omics data can be accomplished by gene set or pathway enrichment methods. A plethora of different tools and algorithms have been developed so far. Among those, the gene set enrichment analysis (GSEA) proved to control both type I and II errors well.

In recent years the call for a combined analysis of multiple omics layer became prominent, giving rise to a few multi-omics enrichment tools. Each of which has its own drawbacks and restrictions regarding its universal application.

Here, we present the `multiGSEA` package aiding to calculate a combined GSEA-based pathway enrichment on multiple omics layer. The package queries 8 different pathway databases and relies on the robust GSEA algorithm for a single-omics enrichment analysis. In a final step, those scores will be combined to create a robust composite multi-omics pathway enrichment measure. `multiGSEA` supports 11 different organisms and includes a comprehensive mapping of transcripts, proteins, and metabolite IDs. It is publicly available under the GPL-3 license at https://github.com/yigbt/multiGSEA and at Bioconductor: https://bioconductor.org/packages/multiGSEA.

## 1 Introduction

When measuring molecular responses to a certain treatment or gaining insights into clinical phenotypes, gene set or pathway enrichment techniques are tools of first choice to infer mechanistic biological information from high-dimensional molecular omics data. Through different statistical techniques, such as overrepresenation analysis (ORA) or gene set enrichment analysis (GSEA), these methods are capable of identifying specific sets of genes or molecular response/signaling pathways that are triggered upon a certain treatment or disease. These sets might represent specific molecular functions, as defined by Gene Ontology (GO) (Gene Ontology Consortium, 2015), biological processes or experimentally derived gene sets which are publicly available in databases such as Reactome (Matthews *et al.*, 2009) or the Molecular Signature Database (MSigDB) Liberzon *et al.* (2011).

Up to date nearly one hundred algorithms have been developed for gene set or pathway enrichment. Each of which has its own strengths and weaknesses. In principle, these methods can be grouped into two distinct classes: 1) pure gene set enrichment, where the algorithms solely focus on a plain list of features and 2) topology-based enrichment, where algorithms include additional information derived from pathway or network databases, e.g., which genes or proteins are directly connected and how they influence each other. There are several comprehensive reviews on this topic available, see for example (Khatri *et al.*, 2012; Nguyen *et al.*, 2019). Besides a plain quality assessment of different enrichment techniques, this review also evaluated the robustness of available methods, i.e., how error-prone these methods are w.r.t. the prediction of either false positive or false negative gene sets or pathways. The popular GSEA method showed a decent quality in terms of rank- and p-value-based pathway enrichment but moreover was the only method not found to produce any false prediction (Nguyen *et al.*, 2019).

Over the last decade, combined analysis of molecular responses through the integration of multiple omics types has become rather frequent, e.g. combining transcriptomics, proteomics, and metabolomics. This is becoming necessary since single-omics analyses will only measure biomolecules of a specific type and will often not even detect its entirety but only a subset thereof. Furthermore, the response time and the life-span of biomolecules varies substantially within and between single omics layers. Thus, only the combined analysis of several molecular layers through multi-omics measurements reliably allows to uncover a significant fraction of cellular effects (Canzler *et al.*, 2020).

There are a few integration tools available that incorporate pathway knowledge to interpret multi-omics datasets like *PaintOmics* (Hernández-de Diego *et al.*, 2018) or *IMPaLA* (Kamburov *et al*., 2011). These methods exhibit several limitations hampering their unrestricted application. *PaintOmics*, for example, is capable of including several different omics layers into its pathway enrichment analysis, but solely relies on pathway definitions from the KEGG database. Furthermore, impacted pathways are determined based on Fisher’s exact test, which was shown to be particularly prone to reporting false positive pathway enrichments (Nguyen *et al.*, 2019). *IMPaLA* on the contrary supports the analysis of a range of different pathway databases but is limited to two different omics input layers, allowing to integrate either transcriptome and metabolome or proteome and metabolome. While *PaintOmics* is applicable to several model organisms (mouse, rat, fruit fly, etc.) *IMPaLA* is restricted to human pathway definitions only.

Here we introduce the `multiGSEA` R package that provides multi-omics-based pathway enrichment employing the robust GSEA algorithm and allowing to use pathway or gene set definitions from several curated databases. In its current version `multiGSEA` is applicable to a combination of transcriptome, proteome, and metabolome data measured in 11 different organisms, including human, mouse, or rat.

## 2 Workflow

In principle, the workflow of the `multiGSEA` package is composed of three essential steps: (i) prepare pathway definitions and omics data (ii) single omics gene set enrichment analysis (iii) combined multi-omics enrichment. These steps are graphically outlined in Figure 1 and described in detail below:

**Figure 1:**
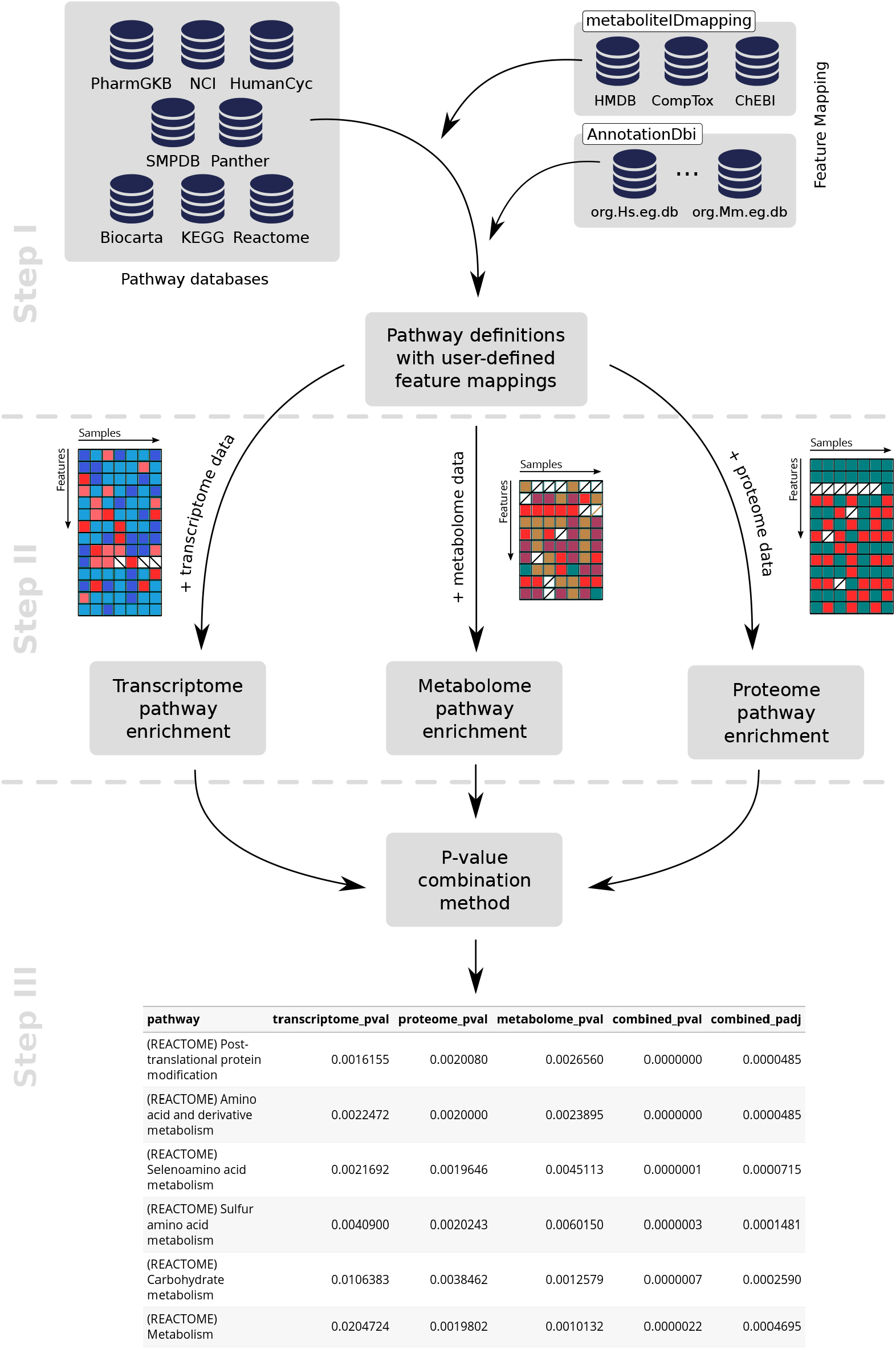
Figure illustrating the workflow of the `multiGSEA` package. In step 1 pathway databases are queried, the features are extracted and mapped to user-defined ID formats. Single omics pathway enrichment using a GSEA-based approach is performed in step 2. P-values from these calculations are combined in step 3 to create a multi-omics pathway enrichment.

### 2.1 Collecting pathway definitions, feature extraction, and mapping

Over the last decades, several pathway databases have been established. Some of which are peer-reviewed and manually curated while others follow a community-based approach to develop and refine pathways. However, often these databases contain their own format in which pathway definitions are provided, making it cumbersome to include multiple databases in an analysis workflow. The ‘graphite‘ ‘R‘ package (Sales *et al.*, 2012) was designed to bridge this gap since it is able to provide pathway definitions from eight publicly available databases – the numbers of currently available human pathway definitions in these databases are listed in parentheses: `KEGG` (311) (Ogata *et al.*, 1999), `Biocarta` (247), `Reactome` (2208) (Matthews *et al*., 2009), `NCI/Nature Pathway Interaction Database` (212) (Schaefer *et al.*, 2009), `HumanCyc` (48682) (Caspi *et al.*, 2010), `Panther` (94) (Mi *et al.*, 2013), `smpdb` (48668) (Jewison *et al.*, 2014), and `PharmGKB` (66) (Whirl-Carrillo *et al.*, 2012). Within the first step of the `multiGSEA` workflow, we make use of the `graphite` package to retrieve pathway definitions from up to eight public databases.

Depending on the database, pathway features (nodes) are encoded with different ID formats. The `KEGG` database, for example, uses Entrez Gene IDs for transcripts and proteins while KEGG Compound IDs are used for metabolites. The `Reactome` database on the contrary stores transcripts and proteins by means of Uniprot identifiers, while ChEBI IDs are used for metabolites. Further metabolite ID formats are CAS numbers and Pubchem IDs. To solve this issue, especially when dealing with multiple pathway databases in a single analysis, we implemented an ID mapping for features of all three supported omics layers. Transcriptomics and proteomics features can be mapped to the following formats: Entrez Gene IDs, Uniprot IDs, Gene Symbols, RefSeq, or Ensembl IDs. The mapping procedure is accomplished by means of the `AnnotationDbi` Bioconductor package (Pagès *et al.*, 2019) and depends on the loaded annotation database such as `org.Hs.eg.db` for human (Carlson, 2019).

Metabolomic features can be mapped to Comptox Dashboard specific IDs (DTXSID, DTXCID), CAS numbers, Pubchem IDs (CID, SID), KEGG Compound IDs, HMDB IDs, or ChEBI IDs. For enhanced usability we encapsulated this comprehensive metabolite mapping data set in a standalone `AnnotationHub` package called `metaboliteIDmapping` (Canzler, 2020). In its current version the package contains more than 1.1 million compounds and was collected and integrated from four different databases: `Comptox Dashboard`^1^ ^2^, `HMDB`^3^, and `ChEBI`^4^.

### 2.2 Gene set enrichment analysis

Measured omics data are necessary for the calculation of gene set enrichment scores. These data have to be loaded for each of the omics layers that have been defined in the previous step of extracting pathway-specific features from external databases. Prior to the enrichment score computation, a differential expression analysis has to be performed such that all omics features have an associated fold change and p-value. The pre-processing step has to be done externally and is not part of the `multiGSEA` package.

In a second step, `multiGSEA` calculates the enrichment score by applying the `fgsea R` package (Korotkevich *et al.*, 2019) on each omics layer individually. The algorithm *GSEA* in its original form was first described to shed light on the mechanistic basis of Type 2 Diabetes mellitus (Mootha *et al.*, 2003). The updated and most commonly applied version was introduced by Subramanian *et al.* (2005) two years later. In brief, measured omics features are utilized for a differential expression testing to derive fold changes and associated p-values. Both values are used to calculate the so-called local statistic, i.e., a ranked gene list based on the direction of the fold-change and the log-transformed p-value. In the following step, GSEA algorithms test whether gene sets accumulate at the top or bottom of those ordered gene vectors. The `fgsea` version used here is an efficient but yet accurate implementation of the prominent *GSEA* algorithm. Its performance is achieved through implementing a cumulative GSEA-statistic calculation allowing to reuse samples between different gene set sizes (Korotkevich *et al.*, 2019).

After the second part of the `multiGSEA` workflow, each downloaded pathway has been assigned fgsea-based enrichment scores, p-values, and adjusted p-values for each omics layer separately.

### 2.3 Combined multi-omics enrichment

To more comprehensively measure a pathway response, `multiGSEA` provides different approaches to compute an aggregated p-value over multiple omics layers. Because no single approach for aggregating p-values performs best under all circumstances, Loughin proposed basic recommendations on which method to use depending on structure and expectation of the problem (Loughin, 2004). If small p-values should be emphasized, Fisher’s method should be chosen. In cases where p-values should be treated equally, Stouffer’s method is preferable. If large p-values should be emphasized, the user should select Edgington’s method. Figure 2 indicates the difference between those three methods.

**Figure 2:**
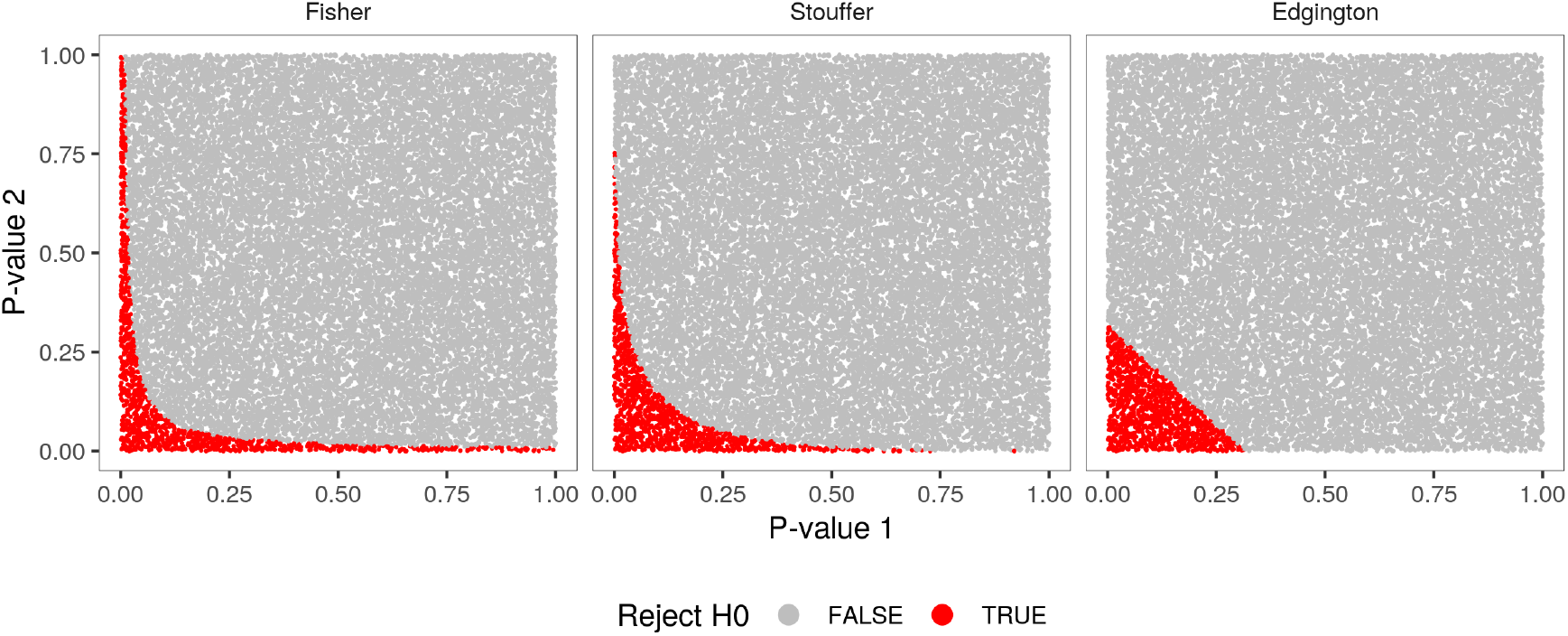
Figure illustrating the difference between Fisher’s combined probability test, Stouffer’s method, and Edgington’s method to aggregate multiple p-values. Two times 25k p-values have been randomly chosen to be subsequently combined by either of those three methods. Combined p-values lower than 0.05 are classified as significant and plotted red.

The first option is the Fisher’s combined probability test, which uses the p-values from *k* independent tests (here up to three omics layer) to calculate a test statistic 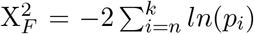. If all of the null hypotheses of the *k* tests are true, the test statistic will follow a X^2^ distribution with 2^*k*^ degrees of freedom (Fisher, 1932). Fisher’s method is asymmetrically sensitive to small p-values which results in a bias for aggregated p-values from multiple studies on the same null hypothesis (Whitlock, 2005).

To circumvent this asymmetry, `multiGSEA` can also apply alternative methods: the Z-transform test and the weighted Z-transform test. The first algorithm is also called Stouffer’s method. Both versions make use of the fact that p-values, ranging from 0 to 1, can be uniquely matched with a value in *Z*, representing a standard normal deviate, and *vice versa.* Each p-value *p_i_* from *k* independent tests (here omics layer) is converted into deviates *Z_i_*, with *Z_i_* = Φ^-1^ (*p_i_*) and Φ being the standard normal cumulative distribution function. Stouffer’s method is defined as:

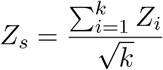

*Z_s_* follows a standard normal distribution if the null hypothesis is true, and thus can be compared to a standard normal distribution to provide a test of the cumulative evidence (Stouffer *et al.*, 1949)

The weighted version of this method is defined by:

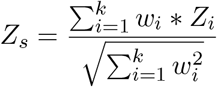

It is still an open debate whether the weighted or unweighted version is preferential. However, it has been reported that if the weighted version is used, optimal results are obtained using weights proportional to the square root of the sample sizes (Lipták, 1958; Zaykin, 2011).

A third alternative method was created by Edgington and relies on untransformed p-values. It was developed to combine probability values through an additive approach (Edgington, 1972):

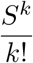

with

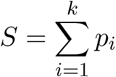

and *k* being the number of individual studies. However, this is a rather conservative estimate resulting in combined probability values that are too high when 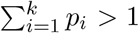. To account for this, correction terms were added to the summation:

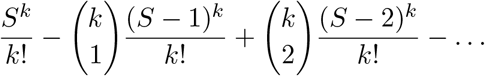

Plus and minus signs alternate and the series continues until the numerator becomes negative. Finally, the result of this progression is compared to a chosen significance level on whether to reject the null hypothesis or not.

A recommendable review on those three (and three additional) aggregation methods was published by Heard and Rubin-Delanchy (2018) alongside some practical advises on how to chose a suitable method.

`multiGSEA` allows to choose from all those different approaches, which are provided through the metap R package (Dewey, 2020).

After combining p-values that were calculated on the same pathway over multiple omics layer by one of the aforementioned methods, `multiGSEA` outputs a ranked table listing those pathways on top that achieved the smallest aggregated p-value.

## 3 Availability

The multiGSEA package is entirely written in R and available under the GPL-3 license. The development version of the package is hosted on GitHub at https://github.com/yigbt/multiGSEA. The package is also part of the Bioconductor project to provide open source software for bioinformatics at https://bioconductor.org/packages/multiGSEA.

For installation we recommend two ways: (i) use the `BiocManager` package from Bioconductor:

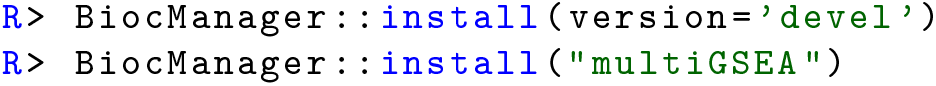

(ii) use the `devtools` library (Wickham *et al.*, 2019) to install directly from our git repository:

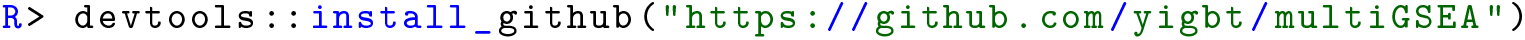

Please visit the repository page to report issues, request features or provide other feedback.

## 4 Example use case

In the following we will briefly cover a use case scenario on human mitochondrial stress data. A comprehensive vignette of the `multiGSEA` package can be found in our git repository or at the Bioconductor package website.

### 4.1 Example data and pathway definitions

At the beginning, the package itself, pathway definitions, and omics data has to be loaded. The datasets in this use case and in the vignette were originally published it is raw form by Quirós *et al.* (2017). Here we make use of processed data that is deposited within the package.

Load the R packages:

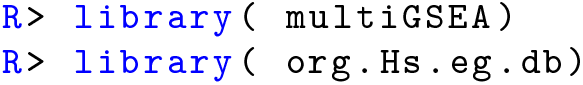

Load the example data sets:

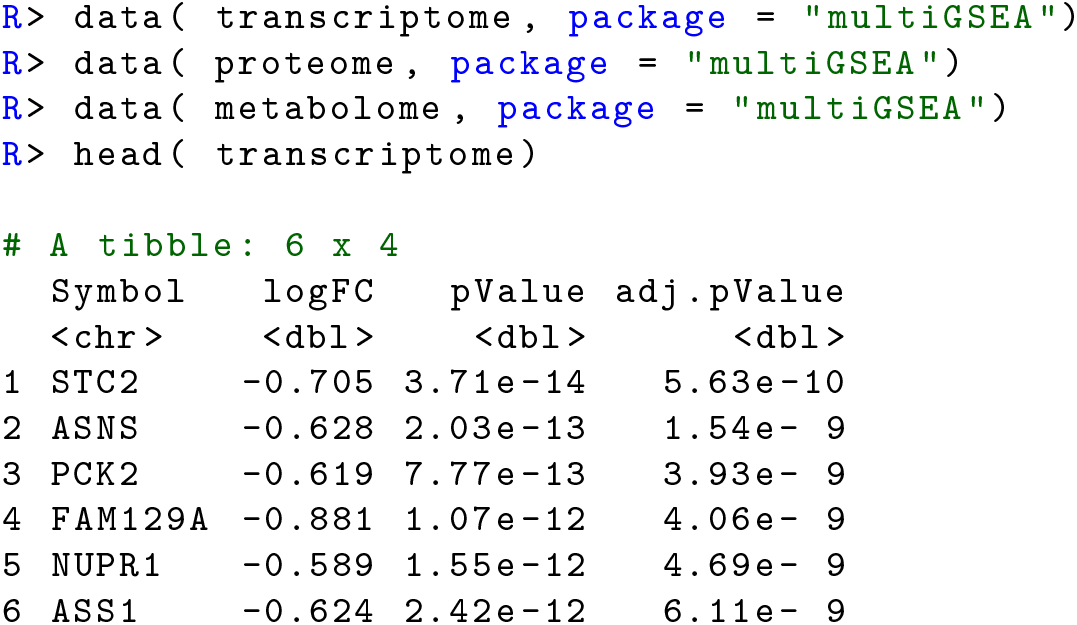

Create a data structure and calculate the pre-ranked feature to speed-up the GSEA procedure later on. `multiGSEA` works with nested lists where each sublist represents an omics layer:

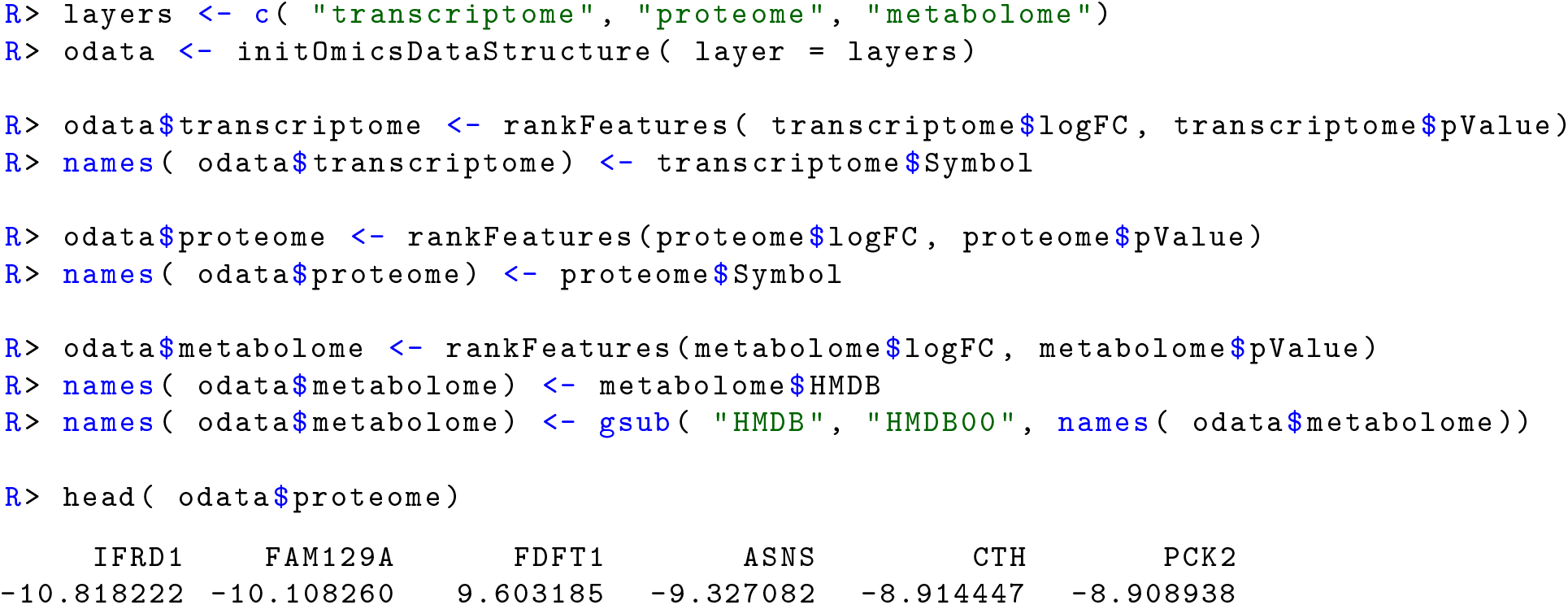

Retrieve pathway definitions and map features to the same ID format as in your omics measurements:

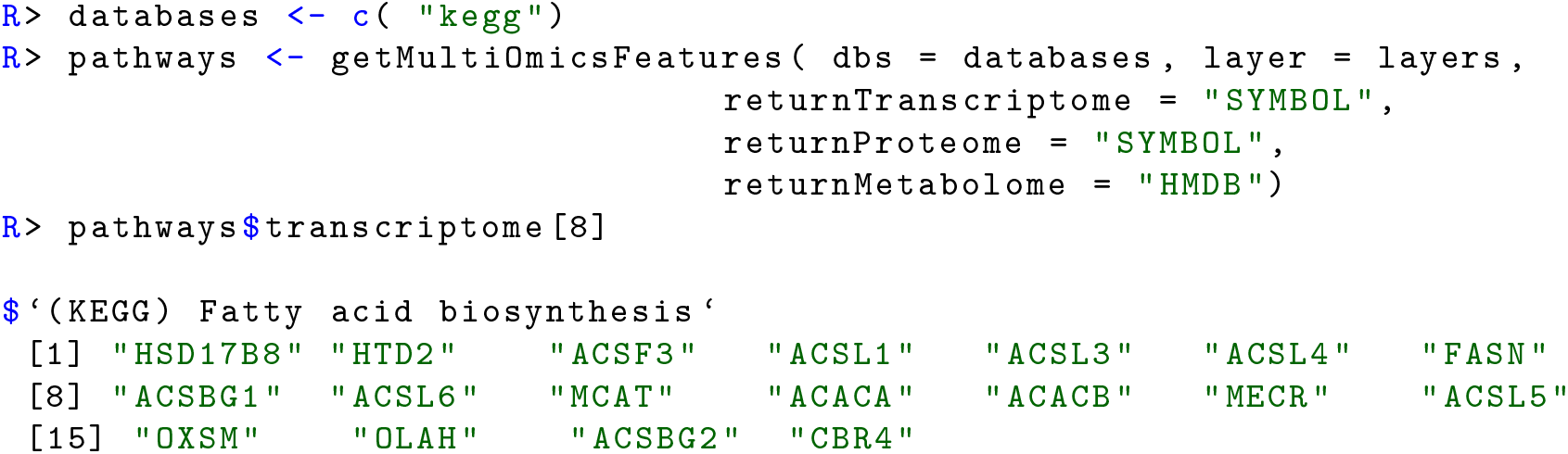

### 4.2 Run pathway enrichment

Now that we have ranked omics features and pre-formatted pathway definitions, we can calculate GSEA-based pathway enrichments for each omics layer separately by means of `multiGSEA`:

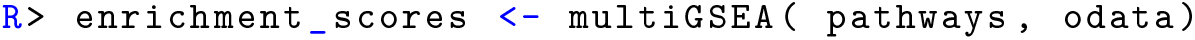

The returned data frame contains nested lists for each analyzed omics layer and each sublist contains the complete gene set enrichment analysis for its respective layer.

### 4.3 Calculate aggregated p-values

For further analysis, the function `extractPvalues` creates a simple data frame where each row represents a pathway and columns represent omics related p-values and adjusted p-values:

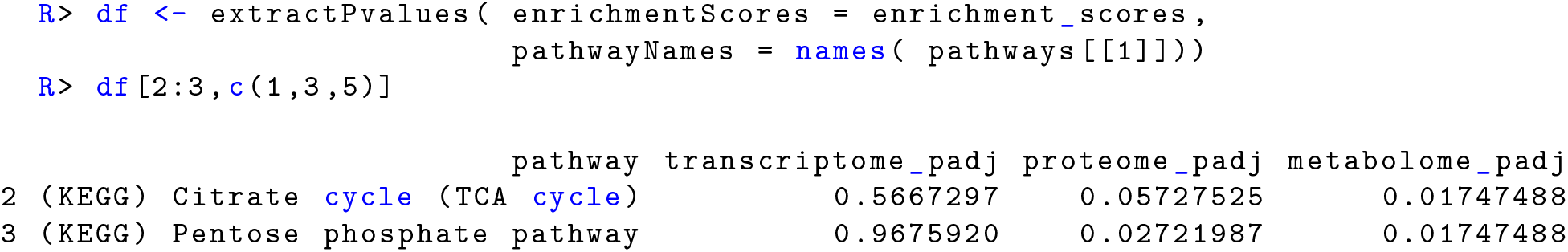

This data structure can then be used to calculate the aggregated p-value and the adjusted p-value:

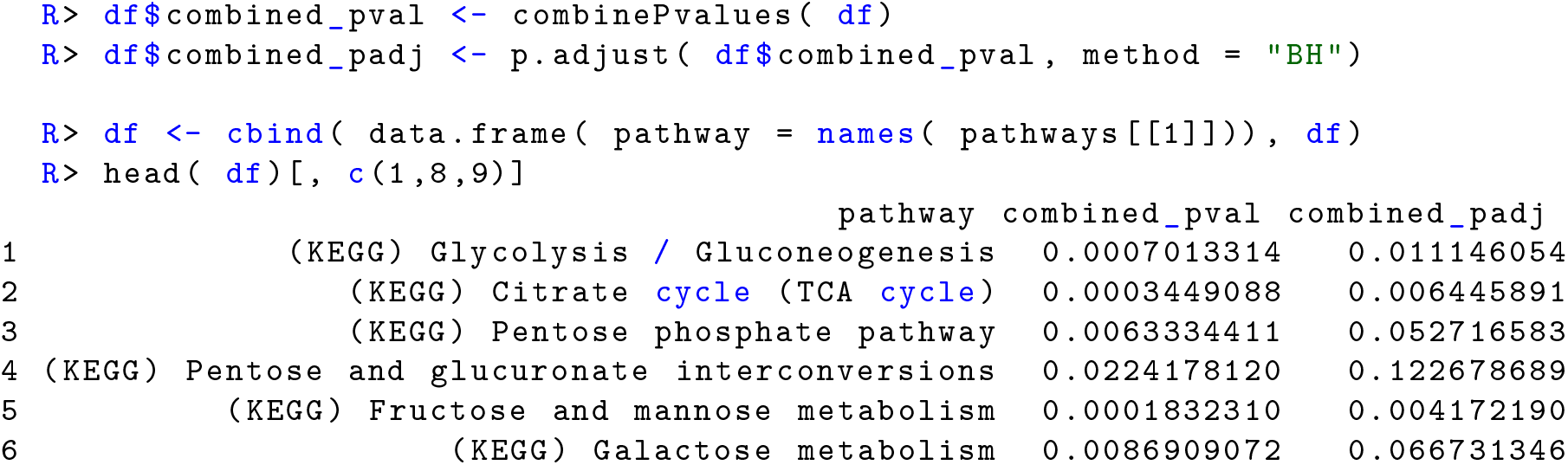

## Acknowledgments

This work was supported in part by the CEFIC Long Range Initiative through funding the project C5 - XomeTox and by the European Union’s Horizon 2020 research and innovation programme under grant agreement No814426 - nanoinformaTIX.

## Footnotes

1 ftp://newftp.epa.gov/COMPTOX/Sustainable\_Chemistry\_Data/Chemistry\_Dashboard/PubChem\_DTXSID\_mapping\_file.txt

2 ftp://newftp.epa.gov/COMPTOX/Sustainable\_Chemistry\_Data/Chemistry\_Dashboard/2019/April/DSSToxSDF.zip

3 http://www.hmdb.ca/system/downloads/current/hmdb\_metabolites.zip

4 ftp://ftp.ebi.ac.uk/pub/databases/chebi/Flat\_file\_tab\_delimited/database\_accession.tsv

## References

Canzler, S. (2020). metaboliteIDmapping. Bioconductor AnnotationHub package version 0.99.4.

Canzler, S. et al. (2020). Prospects and challenges of multi-omics data integration in toxicology. Arch Toxicol, 94(2), 371–388.

Carlson, M. (2019). org.Hs.eg.db: Genome wide annotation for Human. R package version 3.10.0.

Caspi, R. et al. (2010). The MetaCyc database of metabolic pathways and enzymes and the BioCyc collection of pathway/genome databases. Nucleic Acids Res, 38(Database issue), D473–9.

Dewey, M. (2020). metap: meta-analysis of significance values. R package version 1.3.

Edgington, E.S. (1972). An additive method for combining probability values from independent experiments. The Journal of Psychology, 80(2), 351–363.

Fisher, S.R.A. (1932). Statistical Methods for Research Workers - Revised and Enlarged. Edinburgh, London.

Gene Ontology Consortium (2015). Gene Ontology Consortium: going forward. Nucleic Acids Res, 43(Database issue), D1049–56.

Heard, N.A. and Rubin-Delanchy, P. (2018). Choosing between methods of combining P-values. Biometrika, 105(1), 239–246.

Hernández-de Diego, R. et al. (2018). PaintOmics 3: a web resource for the pathway analysis and visualization of multi-omics data. Nucleic Acids Res, 46(W1), W503–W509.

Jewison, T. et al. (2014). SMPDB 2.0: big improvements to the Small Molecule Pathway Database. Nucleic Acids Res, 42(Database issue), D478–84.

Kamburov, A. et al. (2011). Integrated pathway-level analysis of transcriptomics and metabolomics data with impala. Bioinformatics, 27(20), 2917–8.

Khatri, P., Sirota, M. and Butte, A.J. (2012). Ten years of pathway analysis: current approaches and outstanding challenges. PLoS Comput Biol, 8(2), e1002375.

Korotkevich, G., Sukhov, V. and Sergushichev, A. (2019). Fast gene set enrichment analysis. bioRxiv.

Liberzon, A. et al. (2011). Molecular signatures database (MSigDB) 3.0. Bioinformatics, 27(12), 1739–40.

Lipták, T. (1958). On the combination of independent tests. Magyar Tud Akad Mat Kutato Int Kozl, 3, 171–197.

Loughin, T.M. (2004). A systematic comparison of methods for combining p-values from independent tests. Computational statistics & data analysis, 47(3), 467–485.

Matthews, L. et al. (2009). Reactome knowledgebase of human biological pathways and processes. Nucleic Acids Res, 37(Database issue), D619–22.

Mi, H., Muruganujan, A. and Thomas, P.D. (2013). PANTHER in 2013: modeling the evolution of gene function, and other gene attributes, in the context of phylogenetic trees. Nucleic Acids Res, 41 (Database issue), D377–86.

Mootha, V.K. et al. (2003). PGC-1alpha-responsive genes involved in oxidative phosphorylation are coordinately downregulated in human diabetes. Nat Genet, 34(3), 267–73.

Nguyen, T.M. et al. (2019). Identifying significantly impacted pathways: a comprehensive review and assessment. Genome Biol, 20(1), 203.

Ogata, H. et al. (1999). KEGG: Kyoto Encyclopedia of Genes and Genomes. Nucleic Acids Res, 27(1), 29–34.

Pagès, H. et al. (2019). AnnotationDbi: Manipulation of SQLite-based annotations in Bioconductor. R package version 1.48.0.

Quirós, P.M. et al. (2017). Multi-omics analysis identifies ATF4 as a key regulator of the mitochondrial stress response in mammals. J Cell Biol, 216(7), 2027–2045.

Sales, G. et al. (2012). graphite - a Bioconductor package to convert pathway topology to gene network. BMC Bioinformatics, 13, 20.

Schaefer, C.F. et al. (2009). Pid: the pathway interaction database. Nucleic Acids Res, 37(Database issue), D674–9.

Stouffer, S.A. et al. (1949). The american soldier: Adjustment during army life.(studies in social psychology in world war ii), vol. 1.

Subramanian, A. et al. (2005). Gene set enrichment analysis: a knowledge-based approach for interpreting genome-wide expression profiles. Proc Natl Acad Sci U S A, 102(43), 15545–50.

Whirl-Carrillo, M. et al. (2012). Pharmacogenomics knowledge for personalized medicine. Clin Pharmacol Ther, 92(4), 414–7.

Whitlock, M.C. (2005). Combining probability from independent tests: the weighted Z-method is superior to Fisher’s approach. J Evol Biol, 18(5), 1368–73.

Wickham, H., Hester, J. and Chang, W. (2019). devtools: Tools to Make Developing R Packages Easier. R package version 2.2.1.

Zaykin, D.V. (2011). Optimally weighted Z-test is a powerful method for combining probabilities in metaanalysis. J Evol Biol, 24(8), 1836–41.

